# Non-destructive enzymatic deamination enables single molecule long read sequencing for the determination of 5-methylcytosine and 5-hydroxymethylcytosine at single base resolution

**DOI:** 10.1101/2019.12.20.885061

**Authors:** Zhiyi Sun, Romualdas Vaisvila, Bo Yan, Chloe Baum, Lana Saleh, Mala Samaranayake, Shengxi Guan, Nan Dai, Ivan R. Corrêa, Sriharsa Pradhan, Theodore B. Davis, Thomas C. Evans, Laurence M. Ettwiller

## Abstract

The predominant methodology for DNA methylation analysis relies on the chemical deamination by sodium bisulfite of unmodified cytosine to uracil to permit the differential readout of methylated cytosines. Bisulfite treatment damages the DNA leading to fragmentation and loss of long-range methylation information. To overcome this limitation of bisulfite treated DNA we applied a new enzymatic deamination approach, termed EM-seq (Enzymatic Methyl-seq) to long-range sequencing technologies. Our methodology, named LR-EM-seq (Long Range Enzymatic Methyl-seq) preserves the integrity of DNA allowing long-range methylation profiling of 5-mC and 5-hmC over several kilobases of genomic DNA. When applied to known differentially methylated regions (DMR), LR-EM-seq achieves phasing of over 5 kb resulting in broader and better defined DMRs compared to previously reported. This result demonstrated the importance of phasing methylation for biologically relevant questions and the applicability of LR-EM-seq for long range epigenetic analysis at single molecule and single nucleotide resolution.

## Introduction

Long read technologies have been a breakthrough in next generation sequencing for their abilities to phase and resolve variations and repeats over large segments of the human genome [1][2]. Phasing of methylation at single molecule resolution represents a significant advance in addressing the mechanisms and relevance of epigenetic modifications, particularly in repeats, imprinted genes and distant regulatory regions.

Recently, a number of studies have successfully identified cytosine methylation in CpG context with increased accuracy using the ability of the Nanopore sequencer to directly ‘read’ the modification [3][4][5]. Using this method, methylation can be examined over large fragments of genomic DNA. Nonetheless, because the methylation status is not preserved during amplification, only native non-amplified DNA can be used. While enrichment strategies using Cas9 have been applied [6] for targeting specific regions in the genome[7][8], the required starting material is very high and the enrichment is relatively low.

While a number of methodologies have been developed to study cytosine modification [9][10], bisulfite sequencing is still the predominant method used for methylome analysis. Bisulfite sequencing is based on the differential reactivity of cytosine (C) and 5-methylcytosine (5-mC) with sodium bisulfite. Unmodified cytosines are deaminated to uracils (U) and will be read as thymine (T) during sequencing, while 5-mC is unchanged and will be read as “C” [11]. Nonetheless, all bisulfite-based methods introduce DNA strand breaks and results in highly fragmented DNA. This random fragmentation of the deaminated DNA remains the major roadblock to studying epigenetic modifications over large genomic regions using bisulfite sequencing. Indeed, the largest amplicons obtained and sequenced from bisulfite-deaminated DNA does not exceed 1500 bp in length [12].

Recognizing the substantial limitation of bisulfite sequencing in preserving DNA integrity, APOBEC3A cytidine deaminase was used to achieve base resolution sequencing of 5-hydroxymethylcytosine (5-hmC) while avoiding most of the DNA damage [13]. APOBEC3A is a member of the AID/APOBEC (activation-induced cytidine deaminase /apoliprotein B mRNA-editing catalytic polypeptide-like) family of deaminases and has been shown to be critical to immunoglobulin diversification and antiretroviral defense [14]. APOBEC3A preferentially deaminates cytosine and 5-mC resulting in the formation of uracil and thymine, respectively. Because unmodified cytosine and 5-mC are both substrates for APOBEC3A [13], the identification of 5-mC using APOBEC3A alone is currently not possible.

A commercial technology for the determination of 5-mC called EM-seq has become recently available (Materials and Methods). This technology relies on the enzymatic treatment of DNA and thus eliminates the need for bisulfite conversion entirely. In this work, we show that such enzymatic treatment preserve the integrity of the DNA with no detectable evidence of fragmentation or damage. We therefore adapted EM-seq to long read sequencing of amplicon using both PacBio and Nanopore sequencing technologies and extended the technology to both 5-mC and 5-hmC detection. The resulting method, termed herein Long-Read-EM-seq (LR-EM-seq), utilizes the selective enzymatic protection of 5-mC and/or 5-hmC prior to enzymatic deamination by APOBEC3A and large fragment sequencing sample preparation to accurately profile both 5-mC and 5-hmC at base resolution. The preservation of DNA integrity allows the locus-specific amplification of several kb of genomic DNA and the long-range phasing at molecular resolution of 5-mC and 5-hmC. Applied to known differentially methylated regions (DMR) in the mouse genome, LR-EM-seq accurately identifies 5-mC and 5-hmC in more than 5 kb long amplicons allowing the assignment of cytosine modifications to specific alleles.

## Result

### Accurate identification of 5-mC and 5-hmC modification

To enzymatically discriminate epigenetically important cytosine modifications, EM-seq employs the overall strategy depicted in **Fig. 1**. The basic principle of the method consists of selectively modifying epigenetic marks thus protecting those marks from deamination by APOBEC3A. To protect 5-hmC from deamination, 5-hmC is glucosylated with DNA beta-glucosyltransferase (BGT) prior to deamination. This strategy published recently has been shown to effectively discriminate 5-hmC from C and 5-mC [15].

**Figure 1.**
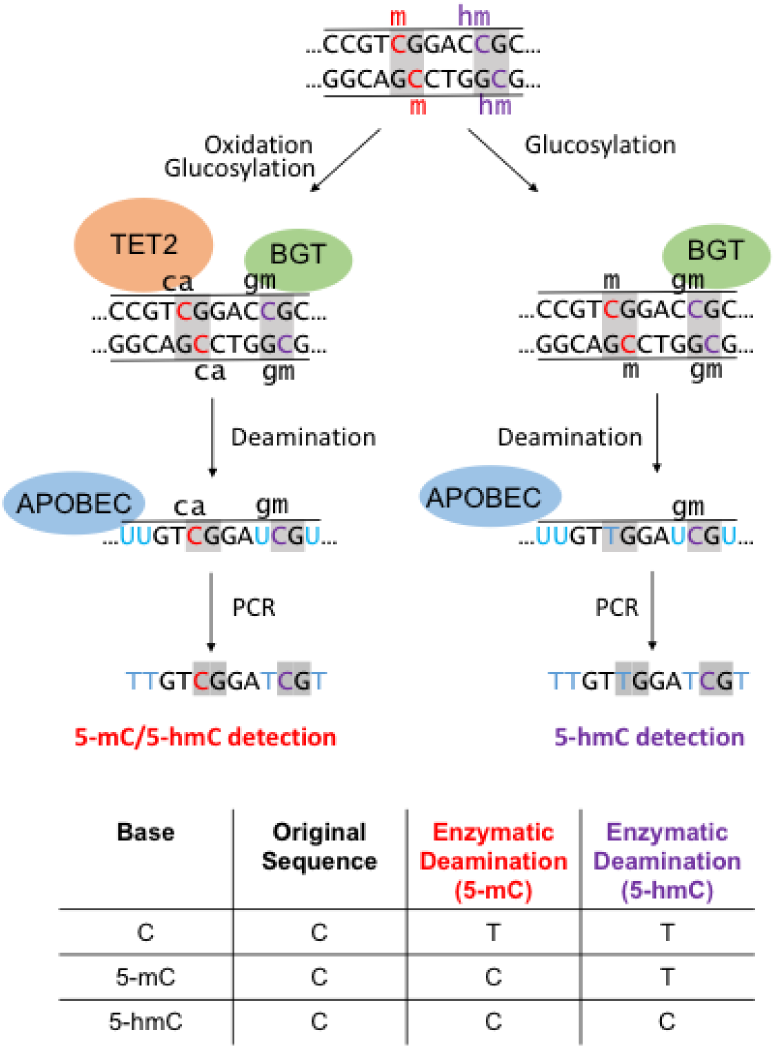
Principle of the EM-seq methodology: genomic DNA can either be treated with TET2 and BGT (left) to protect both 5-mC and 5-hmC, or BGT alone (right) to protect 5-hmC. Subsequent deamination by APOBEC3A followed by PCR amplification allows the distinction between the unprotected substrate (read as T) from the protected cytosine derivatives (read as C).

To discriminate 5-mC from C, the 5-mC needs to be protected from deamination prior to APOBEC3A treatment. 5-mC is converted to glucosylated hydroxymethylcytosine (5-gmC), formylcytosine (5-fC) and carboxycytosine (5-caC) using a combination of 5-mC dioxygenase TET2 and BGT (EM-seq; [16]). All these oxidative products have been shown to be protected from deamination by APOBEC3A including 5-hmC after glucosylation by BGT (EM-seq and [13]).

We independently validated that the strategy described in **Fig. 1a** for the correct identification of 5-mC and compared to Whole-Genome Bisulfite Sequencing (WGBS). While similar accuracy was observed for both methods on CpG sites, enzymatic deamination method showed better accuracy on CpH sites (Supplementary text 1).

To demonstrate that this strategy also results in an accurate identification of 5-hmC, we prepared enzymatic 5-hmC libraries using 50 ng mouse embryonic stem cell (E14) genomic DNA spiked with unmethylated lambda, cytosine methylated XP12 and hydroxymethylated T4gt phage genomic DNAs. Lambda and XP12 control DNA’s were used to measure the deamination rates of APOBEC3A on C and 5-mC, respectively and T4gt DNA was employed to monitor 5-hmC protection from deamination. Using these controls, we calculated the non-conversion rates to be 0.1% for unmodified cytosines; and 1.3% and 1.6% for 5-mCpG and 5-mCH sites, respectively (**Supplementary Table S2**). These non-conversion rates are in line with those reported for TAB-seq and ACE-seq [17][13] (**Supplementary Table S2**). Furthermore, the converted methylated cytosines in XP12 showed no sequence specificity (**Supplementary Fig. 1H**), demonstrating the lack of context bias by APOBEC3A. Importantly, we observed a 98.4% protection rate of 5-hmC by BGT, which is notably higher than that published for TAB-seq (75-92%) (**Supplementary Table S2** and [17]). Thus, our method is expected to have fewer false negative hydroxymethylation calls compared to the widely used TAB-seq method and is in line with the performance of ACE-seq [13]. We also made enzymatic 5-hmC libraries from 1 ng genomic DNA of mouse ES cells. The low input libraries showed similar conversion rates and hydroxymethylation results as the 50 ng libraries (Supplementary Table S5). Both 1 ng and 50 ng libraries gave accurate information on the 5-hmC abundance and deposition at important genomic features, such as epigenetically relevant histone marks, enhancers and transcription factor binding sites (Supplementary text 2 and Supplementary Figs. 1G and 1J).

To measure the range of sensitivity, we shallow-sequenced enzymatically-treated DNA derived from six mouse cell types/ tissues that have been reported to have a wide range of global 5-hmC [18]. We also included a DNMT triple-knockout (TKO) J1 ES cells as a negative control (**Supplementary Table S3**). The average CpG hydroxymethylation level measured after sequencing correlated well with Liquid Chromatography with tandem mass spectrometry (LC-MS/MS) data (r2=0.98, **Supplementary Fig. 1I**). The correlation was linear across a wide range of global 5-hmC levels, demonstrating that the sensitivity is accurate across levels of 5-hmC typically found in mammals [18]. TKO control showed 0.1% of false positive 5-hmC calls indicating an exceptional low non-conversion error rate.

Conducting both enzymatic 5mC and 5hmC sequencing in parallel enabled for the first time the simultaneous investigation of 5-mC and 5-hmC using the same baseline enzymatic reaction (Supplementary Text 2 and Supplementary Figs. 1G and 1J).

### Enzymatic deamination preserves DNA integrity

Having demonstrated the specificity and sensitivity of the enzymatic method towards the identification of 5-mC and 5-hmC, we investigated the ability of the method in preserving the integrity of the DNA, notably for long genomic DNA fragments.

To directly assess the DNA integrity after bisulfite and enzymatic treatments, we separately addressed the loss of amplifiable material for small and large target sizes. For small target sizes (less than 1 kb) we used real-time PCR-based assay to quantify the amount of intact material. With a target size of 809 bp, the relative amount damage-free amplifiable template in the bisulfite treated samples was 94% reduced compared to enzymatic deamination (**Fig. 2a**), which is consistent with previously reported degradation rates of bisulfite method [19].

**Figure 2.**
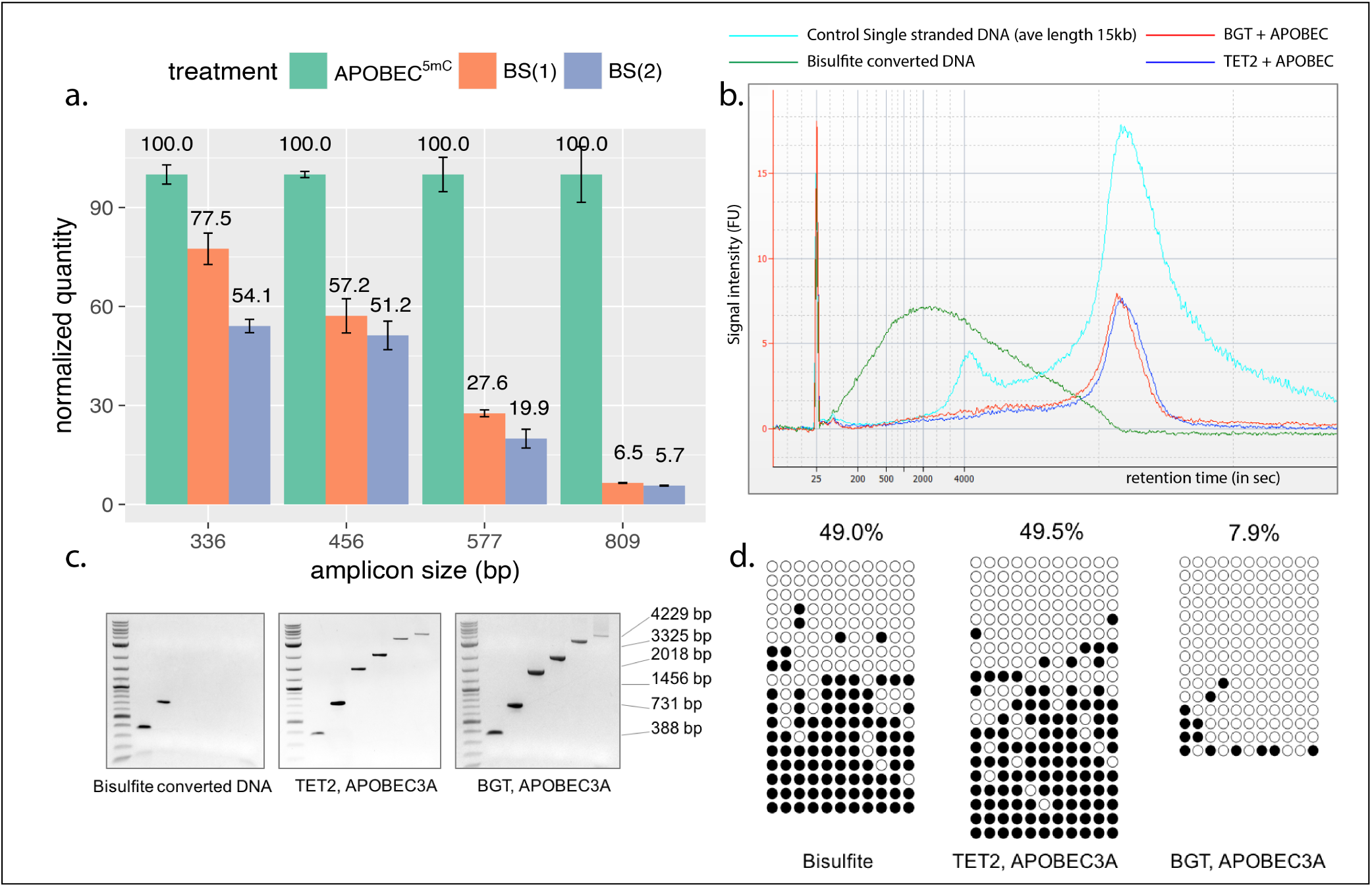
Enzymatic deamination preserves the integrity of the DNA: **a.** qPCR results show the quantities of undamaged amplifiable DNA templates of different sizes after the enzymatic deamination (green) and bisulfite treatments (orange and blue). All quantifications are normalized to the values obtained for the enzymatic deamination experiments. **b.** Agilent 2100 Bioanalyzer trace on RNA 6000 pico chip comparing equal amounts of mouse E14 genomic DNA sheared to an average of 15 kb and treated with sodium bisulfite (green), BGT and APOBEC3A (red), or TET2 and APOBEC3A (blue) over the control ssDNA (magenta). Bisulfite treatment fragmented the DNA to an average of 800 bp, while enzymatically treated DNA show no notable size differences compared to control DNA. **c.** Agarose gel images of end-point PCR of six amplicons ranging from 388–4229 bp illustrating upper amplicon size limit for sodium bisulfite, TET2 and APOBEC3A, or BGT and APOBEC3A treated E14 genomic DNA. d. 731 bp amplicons from the agarose gels showed in (c) were cloned, sequenced and the methylation status determined for bisulfite (left panel), enzymatically converted for 5-mC (center panel) and 5-hmC (right panel) E14 genomic DNA. Open and closed circles indicate unmethylated and methylated, respectively.

Next, we fragmented the genomic DNA to an average of 15 kb and profiled the fragment size distribution before and after enzymatic and bisulfite treatment. Expectedly, the average fragment size after bisulfite treatment dropped substantially from 15 kb to only 0.8 kb (**Fig. 2b**). Sharply contrasting to bisulfite conversion, enzymatic treatment of the same starting amount of DNA conserved the original 15 kb average size profile observed in the control DNA (**Fig. 2b**). This result demonstrates the enzymatic deamination method does not introduce strand breaks even in the case of large DNA fragments.

Lastly, to assess the ability to amplify the DNA material described above, we designed six pairs of primers with a range of predicted amplicon sizes ranging from 388 to 4226 bp. In line with the DNA integrity assessment data, amplification products from bisulfite treated DNA were only detected up to 731 bp. In contrast, all amplicon sizes were amplifiable after enzymatic deamination for both 5-mC and 5-hmC detection (**Fig. 2c**). Sanger sequencing of the 731 bp amplicons showed a nearly identical methylation profile for both enzymatic and chemical deamination methods (**Fig. 2d**), confirming that enzymatic deamination method can provide the same accuracy of methylation detection as bisulfite treatment without damaging the DNA.

### Methylome and hydroxymethylome phasing using long-read sequencing

Preserving the integrity of genomic DNA after enzymatic deamination offers the unique opportunity to study long range epigenetic marks at single base and molecule resolution beyond the reported 1.5kb region achieved using SMRT-BS [12]. As a proof of principle, we apply LR-EM-seq to a 5378 bp region of the mouse genome using DNA derived from ES cells. Two control DNA consisting of CpG methylated pUC19, and unmethylated lambda DNA were spiked to the mouse genomic DNA prior to any enzymatic reactions. For 5-hmC detection, an additional control consisting of T4gt genomic DNA was included to the spike-ins in order to monitor the 5-hmC protection rate. Following enzymatic treatment, 5378 bp mouse amplicon, 3233 bp lambda amplicon, 1774 bp pUC19 amplicon and 5349 bp T4gt amplicon (for 5-hmC detection) were obtained and sequenced using all 3 major sequencing platforms: Oxford Nanopore, PacBio and Illumina. In the case of Illumina sequencing, the amplicons were fragmented to a mean of 600 bp for compatibility with short read sequencing.

Using Illumina data for 5-mC detection, lambda amplicon shows non-conversion error rates of 0.1% whereas the CpG methylated pUC19 amplicon shows 97.4% 5-mC protection rate by the TET2/BGT enzymes. For 5-hmC detection, the non-conversion error rate of cytosine is 0.1%, the non-conversion error rate of 5-mC is 0.6%, and the protection rate of 5-hmC measured using the T4tg amplicon is 99.4%. These values are consistent with our WGBS data obtained with short fragments. In these experiments, the enzymatic treatment and amplification was done on an unfragmented genomic template demonstrating that the enzymatic deamination method is applicable for the long DNA fragments as effectively as for the short DNA material of the WGBS applications.

PacBio sequencing gave very similar estimates to the Illumina results, while Nanopore sequencing generated slightly higher error rates, presumably because of the intrinsic higher error rate of Nanopore sequencing (**Table 1**). At single base resolution, both methylation and hydroxymethylation levels of CpG sites recorded from different sequencing platforms are highly correlated and the modification profiles across the entire region are in agreement (**Figs. 3a, b**). We also compared 5-hmC results with publically available datasets derived from Pvu-Seal-seq [**?**] and TAB-seq [**?**] from the same cell line and found consistent results with our data (**Supplementary Fig. 3**). These results suggest that the LR-EM-seq method is compatible with all the major sequencing platforms and produces accurate results for both 5-mC and 5-hmC. Most significantly, at single-molecule resolution LR-EM-seq coupled with long-read sequencing technologies (PacBio and Nanopore) can provide complete 5-mC and 5-hmC information of entire molecules (**Fig. 3c**) and thus make it possible to study the relationships between distant cytosine sites as well as between individual molecules.

**Table 1:** Percentage of 5-mC or 5-hmC in CpG and CpH contexts (with H = A or T or C) in amplicons derived from Lambda (unmethylated cytosines), pUC19 (CpG methylation), T4gt (hydroxymethylated cytosines) and mouse genomic DNA measured using three sequencing platforms (Illumina, PacBio and Oxford Nanopore).

|  |  | 5-mC |  |  | 5-hmC |  |  |
| --- | --- | --- | --- | --- | --- | --- | --- |
|  |  | Illumina | Pacbio | Nanopore | Illumina | Pacbio | Nanopore |
| Lambda | CpG | 0.1 | 0.1 | 1.1 | 0.1 | 0.1 | 1.1 |
|  | CpH | 0.1 | 0.1 | 1.6 | 0.1 | 0.1 | 1.7 |
| pUC19 | CpG | 97.4 | 97.6 | 90.2 | 0.6 | 0.9 | 2.1 |
|  | CpH | 2 | 0.4 | 2 | 0.2 | 0.6 | 1.7 |
| T4gt | CpG | NA | NA | NA | 99.9 | 99.8 | 87.6 |
|  | CpH | NA | NA | NA | 99.4 | 98.5 | 91.0 |
| mouse | CpG | 73.2 | 76.2 | 67.3 | 5.5 | 5.9 | 5.8 |
|  | CpH | 0.4 | 0.5 | 1.9 | 0.1 | 0.1 | 1.2 |

**Figure 3.**
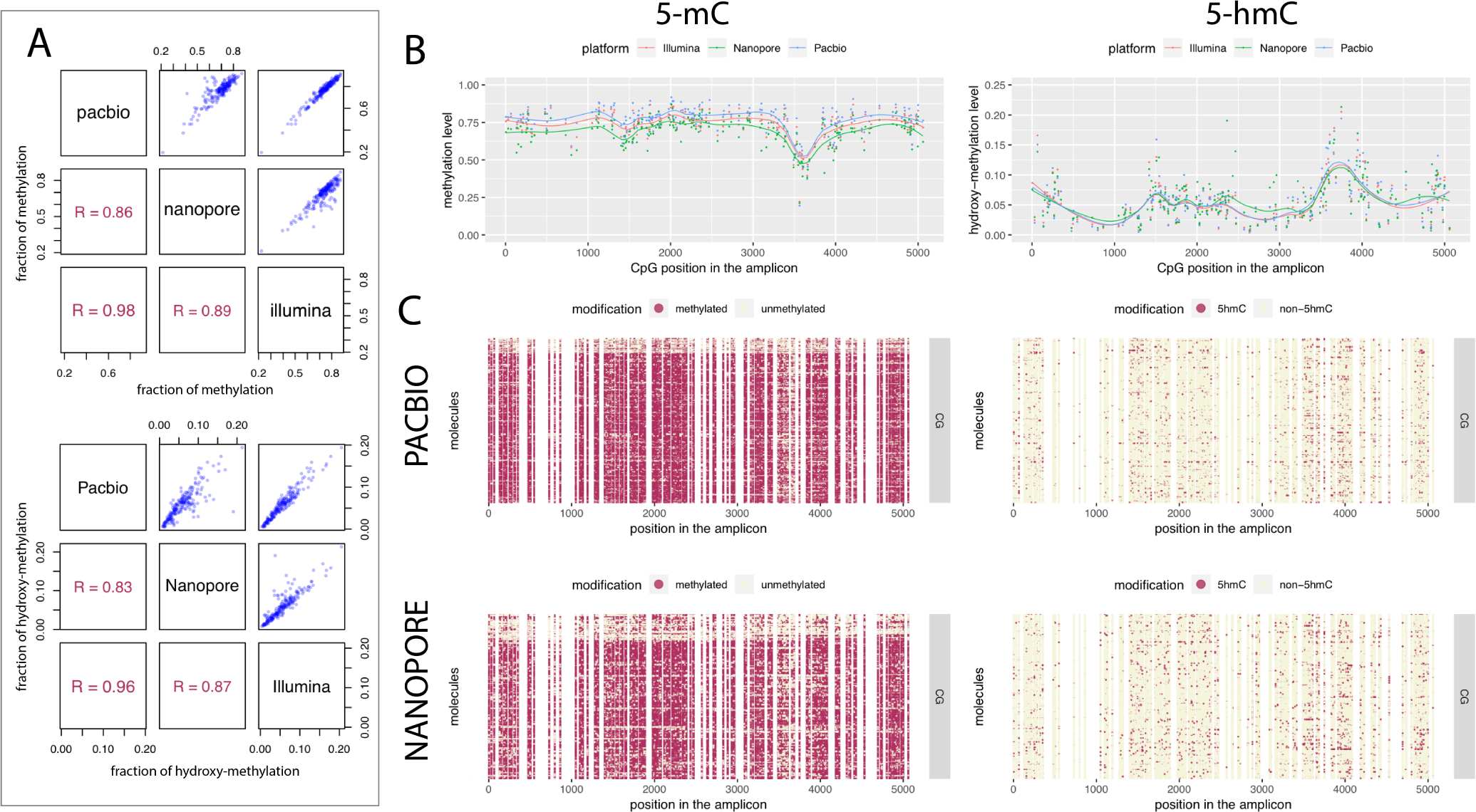
5-mC and 5-hmC phasing using long read sequencing. **a.** Scatter plots and Pearson correlations of calculated methylation (left) and hydroxymethylation (right) levels of all CpG sites within the 5,378 bp region between the 3 sequencing platforms: PacBio, Nanopore and Illumina. **b.** Dot plots showing methylation (left) and hydroxymethylation (right) levels of individual CpG sites within the 5,378 bp region calculated by the LR-EM-seq method using 3 major sequencing platforms: Illumina (red), Nanopore (green) and PacBio (blue). The fitted lines are drawn using the LOESS method. **c.** Single-base single-molecule cytosine modification maps of the 5,378 bp region from the mouse E14 genome generated by the LR-EM-seq method coupled with PacBio SMRT sequencing (top) and Nanopore sequencing (bottom). Methylated (left) and hydroxymethylated CpG sites are depicted by maroon dots and unmodified CpG sites are depicted by beige dots.

Next, we applied LR-EM-seq using SMRT sequencing to a 4614 bp region (chr7: 135829567-135834180, mm9) containing a known 367 bp DMR upstream of a previously described imprinted gene Inpp5f.v2 in mouse brain [20]. Based on the methylation call in CpC and CpT context, the overall conversion rate of the APOBEC3A treated DNA was 99.8% (**Supplementary Table S5**), which is consistent with the performance of LR-EM-seq and corresponds to about 10-fold lower non-conversion rate compared to previously published SMRT-BS sequencing (97.3%) [12]. The methylation profiles showed a clear segregated pattern at the known DMR confirming the differential methylation of this region (**Fig. 4a**). Phasing the entire 4614 bp region allowed a precisely delimitation of the boundary of the DMR at molecule resolution. As a result we report a more than two-fold increase in the size of the reported DMR region from 367 bp to 1 kb (**Figs. 4a,b**). Moreover, when correlating long range methylation pattern, we found two sub-domains flanking both sides of the newly identify DMR (**Fig. 4b**). This suggests the occurrence of differentially methylated domains, whose methylation patterns do not completely follow the core DMR but are correlated with it. Whether such domains are derived from the core DMR under relaxed pressure, serve as a buffer between DMRs and non-DMRs, or indicate independent trans-acting transcription factor binding sites, awaits further investigation. Another important observation is that the CpA, but not CpY cytosine methylation is specifically missing from the extended DMR and mCpA displayed an oscillating pattern around the DMR (Supplementary **Figs. 4a,b**). This suggests that CpA could be the true methylation motif, and high-level chromatin structure, for example nucleosome positioning, may play a role in the deposition of DNA modification and gene regulation near DMR.

**Figure 4.**
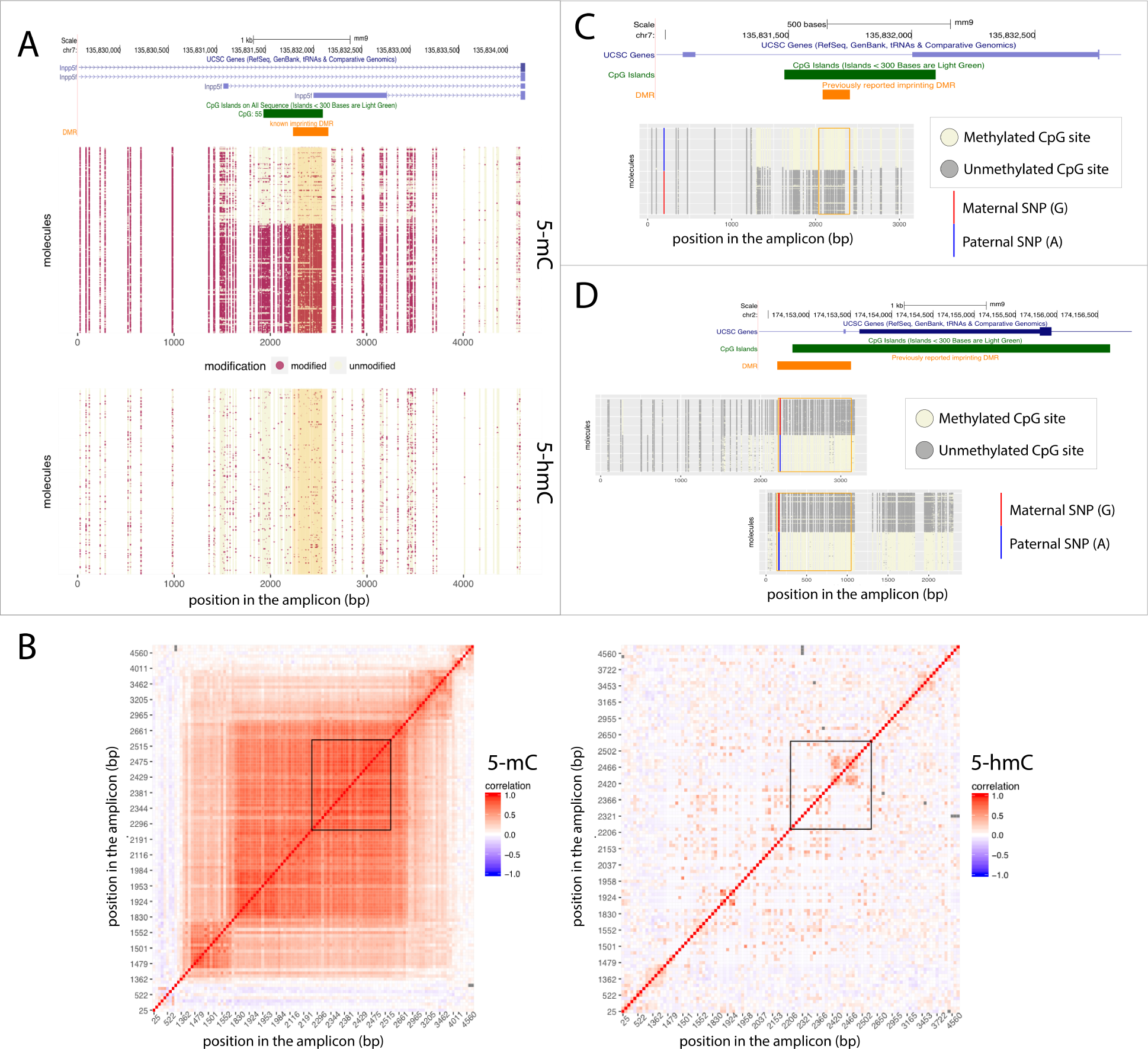
Phasing of DNA modifications and genetic variants by LR-EM-seq. **a.** 5-mC (middle panel) and 5-hmC (bottom panel) state (beige: unmodified; marron: modified) of individual CpG sites at single molecule resolution of the 4.6 kb region overlapping the promoter of the imprinted Inpp5f.v2 gene (top panel) which includes a previous determined DMR (orange box). The shaded area in the dot plots corresponds to the known DMR. **b.** Correlations of CpG modification state (left panel: 5-mC, right panel: 5-hmC). Each cell in the matrix represents the correlation of 2 CpG sites’ modification state across all the studied molecules. The correlation strength is depicted by color (red: correlation=1, blue: correlation=-1). The known DMR is indicated by a black box. **c.** Phasing of 5-mC and single nucleotide polymorphism (SNP) of a 3.1 kb region in the imprinted Inpp5f.v2 gene promoter of the mouse cortex brain from a F1 offspring of a cross between two inbred mouse strains (129×1/SvJ male and Cast/EiJ female). Methylation state of individual CpG sites at single molecule level were denoted by either a beige dot (unmodified) or a red dot (methylated). A SNP near the 5’ end of the region was highlighted with “A” for paternal allele and “G” for maternal allele. The orange box denotes a previously identified imprinted DMR. **d.** Phasing of 5-mC and SNP in the imprinted Gnas1a gene promoter of the mouse cortex brain from a F1 offspring of a cross between two inbred mouse strains (129X1/SvJ male and Cast/EiJ female). Methylation states of individual CpG sites at single molecule level were denoted by either a beige dot (unmodified) or a red dot (methylated). A SNP was highlighted with “A” for paternal allele and “G” for maternal allele. The orange box denotes a previously identified imprinted DMR.

We also successfully phased 5-hmC in the same region. However, we did not observe any significant segregation pattern of hydroxymethylation (**Figs. 4a,b**). At the population level, the average CpG hydroxymethylation abundance significantly decreased at the DMR and generally followed the trend of 5-mC across the entire region (**Supplementary Fig. 4c**), implying that in this region, the hydroxymethylation level may be largely determined by the substrate availability.

We then used LR-EM-seq to validate previously reported allele-specific DMRs in the mouse genome of two inbred mouse strains, 129X1/SvJ (129) and Cast/EiJ (Cast) [21]. For all of the 4 investigated regions (H13, Inpp5f, Gnas1a and Peg12), we observed a segregation of two distinct populations of molecules according to their methylation status i.e., hypermethylated vs. hypomethylated (**Supplementary Fig. 4d**) confirming the existence of a DMR. Moreover, all the DMRs are hundreds of bp to several kb larger than the reported ones with long read sequencing providing precise boundaries (**Supplementary Fig. 4d**).

Next, we used LR-EM-seq to phase both SNP variants and DNA methylation simultaneously to further confirm the allele specificity of the observed DMRs near the imprinted genes. To acquire a large number of heterozygous SNPs, we performed crosses between two inbred mouse strains 129X1/SvJ (129) and Cast/EiJ (Cast) as described in [21]) and use LR-EM-seq to sequence two of the DMR loci (Inpp5f and Gnas1a) in the F1 genome. We identified reliable heterozygous SNPs in both of the amplified DNA fragments. In the case of Inpp5f the heterozygous SNP is almost 2 kb upstream of the DMR (**Fig. 4c**). In both cases, the methylation pattern segregates perfectly with the heterozygous SNP (**Figs. 4c,d**).

The ability to obtain large amplicons greatly expands the genomic ranges that are amenable to phasing of sequence variation with epigenetic information, thus making LR-EM-seq a convenient and promising technology for the identification of allele-specific methylation.

## Conclusion

In this study, we provided compelling evidence for the benefits of enzymatic deamination to identify both 5-mC and 5-hmC using long-read sequencing technology. Importantly, the converted genomic DNA can be amplified and the information regarding the methylation status is preserved allowing for locus-specific interrogation of methylation on low amount of starting material.

Adapting EM-seq to long-read sequencing workflow, termed LR-EM-seq, surpasses bisulfite deamination, notably in producing deaminated DNA without detectable damage. These advances eliminate the foremost roadblocks encountered using bisulfite sequencing for decades. As we have demonstrated in this study, longer DNA material enabled by LR-EM-seq enables the study of the combinatorial effect of methylation over large regions at single-molecule resolution. We also demonstrated the importance and advantage of phasing methylation on long DNA fragments on the study of imprinted genes by identifying much larger DMR regions than previously observed. These previous studies using short-read sequencing have relied on statistical methods to acquire methylation haplotype information and consequently is prone to inaccurate calls. By phasing variation with methylation on a single long read, LR-EM-seq is more accurate and expands the fraction of the genome that can be epigenetically haplotyped.

Phasing methylation has numbers of additional applications, notably in cancer detection where the combinatorial methylation status of several CpG at single molecule resolution is expected to be a much more powerful determinant of tumorigenicity compared to an average methylation level. Combined with other genomic information, phasing methylation relative to variants or epigenetic markers offers an exciting prospective empowered by LR-EM-seq.

Lastly, the ability to amplify longer amplicon provides greater flexibility in primer design, notably in encompassing or avoiding repeats or challenging to amplified regions. These results in larger covered genomic regions with less amplicons.

## Materials and Methods

### E14 Embryonic Stem (ES) Cell culture

ES cells were cultured as previously described [22]. Briefly, cells were grown in GMEM media (Thermo Fisher Scientific, Waltham, MA, USA) containing 10% FBS (Gemcell), 1% non-essential amino acids (NEAA) (Hyclone), 1% sodium pyruvate (Thermo Fisher Scientific, Waltham, MA, USA), 50 µM β-mercaptoethanol (Sigma, St. Louis, MO, USA), and 1 LIF (Millipore, Burlington, MA, USA). To maintain the undifferentiated state, ES cells were grown on 0.1% gelatin-coated culture dishes (Stem Cell Technologies, Vancouver, BC, Canada).

### Genomic DNAs

Mouse genomic DNA from brain, spleen, heart, and liver tissues were obtained from BioChain (Hayward, CA, USA), mouse NIH/3T3 and human Jurkat DNA were from NEB (Ipswich, MA, USA). E14 genomic DNA was extracted with a DNeasy Blood and Tissue Kit (QIAGEN, Hilden, Germany). Genomic DNA from DNMT TKO J1 ES cells were obtained from Dr. Yi Zhang

### Control DNAs

Fully CpG methylated pUC19 DNA was acquired by incubating at 37C for 2 h, 3 µg of dam-dcm- plasmid DNA in a 50 µl reaction containing 20 U of M.SssI methylase (NEB, Ipswich MA, USA), 1X NEBuffer 2, and freshly prepared 160 µM S-adenosylmethionine, followed by heat inactivation of the enzyme at 65C for 20 min and SPRI beads purification. LC-MS analysis was used to verify completeness of methylation status. T4gt (amC87(42-), amE51(56-), NB5060(ΔrllB-denB-ac), unf 39(alc) and XP12 phage genomic DNA’s were extracted as described in (Sambrook, Joseph, Edward F. Fritsch, and Tom Maniatis. Molecular cloning: a laboratory manual. No. Ed. 2. Cold spring harbor laboratory press, 1989.) 5-mC free Lambda genomic DNA was purchased from Promega (Madison, WI, USA).

### 5-mC and 5-hmC phasing of the 5.4Kb mouse genomic region using LR-EM-seq

#### Enzymatic deamination for 5-mC detection

For 5-mC detection, 200 ng of mouse E14 genomic DNA was mixed with 10 ng unmethylated lambda DNA, 10 ng of XP12 phage DNA, and 1 ng of CpG methylated pUC19 DNA, then incubated with 16 µg of TET2 enzyme (EM-seq component E7130A, NEB, Ipswich MA, USA) for 30 min at 37°C, in 50 µl 1x reaction buffer(EM-seq TET2 Reaction Buffer (reconstituted), E7128A and E7131A diluted) followed by a 30 min incubation with 20 U of T4 BGT (EM-seq component E7129A, NEB, Ipswich, MA, USA) in the same buffer at 37°C. Oxidized genomic DNA was incubated additional 30 min with 0.8 U of Proteinase K (NEB, Ipswich, MA, USA) at 37°C, and subsequently purified with a Genomic DNA Clean Concentrator (Zymo Research, Irvine, CA, USA). Purified DNA was then denaturated at 90°C in presence of 29% of formamide, for 10 minutes, and deaminated with 100 U of APOBEC3A (EM-seq components E7133AA and E7134AA NEB, Ipswich, MA, USA) in 100 µl reaction volume for 3 h. 3 µl of deaminated genomic DNA and control DNA’s were used without further purification for PCR amplifications with Phusion U Hot Start DNA Polymerase (Thermo Fisher Scientific, Waltham, MA, USA) using primer pairs listed in Supplementary Table 6. For some amplicons (Supplementary Table 6, 8-11 primer pairs), we used Enzymatic EM-seq Conversion Module (NEB, E7125) for 5-mC detection

#### Enzymatic deamination for 5-hmC detection

For 5-hmC detection, 200 ng of mouse E14 genomic DNA was mixed with 10 ng unmethylated lambda DNA, 10 ng of T4gt phage DNA, and 1 ng of CpG methylated pUC19 DNA, then incubated with 20 U of BGT (NEB, Ipswich, MA, USA) in 1X NEBuffer 2 for 2 h at 37°C. Glucosylated genomic DNA was incubated additional 30 min with 0.8 U of Proteinase K (NEB, Ipswich, MA, USA) at 37°C, and purified with a Genomic DNA Clean Concentrator kit (Zymo Research, Irvine, CA, USA). Purified D NA w as t hen d enaturated at 90°C in presence of 29% of formamide, for 10 minutes, and deaminated with 100 U of APOBEC3A (EM-seq components E7133AA and E7134AA) in 100 µl reaction volume for 16 h. 3 µl of deaminated genomic DNA and control DNA’s were used without further purification for PCR amplifications with Phusion U Hot Start DNA Polymerase (Thermo Fisher Scientific, Waltham, MA, USA) using primer pairs listed in Supplementary Table 5.

#### Illumina sequencing of the enzymatic deaminated amplicons

5-mC and 5-hmC amplicons were pooled with the control amplicons respectively. 50 ng of each amplicon pools were fragmented to an average size of 600 bp using the Covaris S2 instrument in 50 µl of 0.1x TE buffer. Sonicated DNA was used to construct libraries with a NEBNext Ultra DNA Library Prep Kit (NEB, Ipswich, MA, USA), and sequenced on an Illumina MiSeq instrument.

#### Nanopore sequencing of the enzymatic deaminated amplicons

The 1D Native barcoding genomic DNA kit (EXP-NBD103 and SQK-LSK108 kits, Oxford Nanopore Technologies, Oxford, UK) was used for library preparation. 5-mC and 5-hmC amplicons were pooled with the control amplicons and were barcoded to allow for multiplexing and sequencing on the same flow cell. For each sample, end-repair and dA-tailing was performed using the NEBNext Ultra II End repair/dA-Tailing module (E7546, NEB, Ipswich, MA, USA), following Oxford Nanopore protocol, except that the incubation time was increased to 20 min. After clean-up following the end-repair step as described in the protocol, the samples were quantified using Qubit fluorometer (Thermo Fisher Scientific, Waltham MA, USA). 500 ng of each pool were ligated to a single barcode (EXP-NBD103 kit, Oxford Nanopore Technologies) following Nanopore protocol, except the ligation time was increased to 30 min. The samples were quantified with Qubit fluorometer (Thermo Fisher Scientific, Waltham MA, USA) and pooled in equimolar amounts to produce a final amount of 500 ng. The Nanopore adapter ligation was performed according to the manufacturer’s protocol. The library was sequenced for a total of 11h on a MinION (Oxford Nanopore Technologies, Oxford, UK) using a FLO-MIN106 Rev D flow cell. Raw fast5 data were generated using MinKNOW version 18.12 and base called using Guppy base caller Version 2.1.3.

#### Single Molecule Real Time (SMRT) sequencing of enzymatic deaminated amplicons

5-mC and 5-hmC amplicons were pooled with the control amplicons and the amplicon pool (400 ng) were ligated to SMRT bell adapters (Pacific Biosciences, Menlo Park, CA, USA) using T4 DNA ligase (NEB, Ipswich, MA, USA), and the incompletely ligated amplicons were removed using Exonuclease III (NEB, Ipswich, MA, USA) and Exonuclease VII (NEB, Ipswich, MA, USA). Then the purified SMRT bell libraries were sequenced on PacBio Sequel platform following manufacturer’s protocols for polymerase binding (Sequencing primer v3 and Sequel binding Kit 2.1, Pacific Biosciences, Menlo Park, CA, USA) and sequencing (Sequel sequencing kit 2.1, Pacific Biosciences, Menlo Park, CA, USA). One SMRT cell (SMRT Cell 1M v2, Pacific Biosciences, Menlo Park, CA, USA) was used for each library with 600 min movie. Circular Consensus Sequences (CCS) were extracted from the raw movie data and converted into fastq file using SMRT Link (version 6.0.0.47841) CCS protocol.

### 5-mC and 5-hmC phasing of mouse DMRs using LR-EM-seq

#### Mice

Three different mice strains are used for this project: 1, Cast/EiJ (Cast); 2, 129×1/SvJ (129); and 3, The F1 offspring of Cast (female) X 129 (male). The crosses of Cast and 129 mice were performed by the Jackson Laboratory (Bar Harbor, ME, USA). The frontal cortex samples of the male mice F1 offspring and a male mouse of each parental strain were collected at 8 to 10 weeks at the Jackson Laboratory and were shipped to investigator at NEB on dry ice (compliant with the provisions of the Public Health Service Policy on Humane Care and Use of Laboratory Animals).

#### Genomic DNA extraction and purification

The genomic DNA was extracted from 10 mg frozen brain cortex samples using NEB Monarch genomic DNA® purification kit (NEB, Ipswich, MA, USA). Four microliters of RNase A (100mg/mL) have been added to the tissue lysate (and incubated 5min at room temperature) in both protocols to prevent the inhibition of APOBEC3A by RNA during the deamination process. The extracted genomic DNA were purified again using AMPure XP beads.

#### Preparation of LR-APOBEC-seq long amplicons

200 ng purified genomic DNA was glucosylated by incubating with 20U of BGT (NEB, Ipswich MA, USA) for 2 h at 37°C (for 5-hmC detection). Glucosylated genomic DNA was incubated additional 30 min with 0.8U of Proteinase K at 37°C, and subsequently purified with Genomic DNA Clean Concentrator kit (Zymo Research, Irvine, CA, USA Research). For 5-mC detection, mouse brain genomic DNA (200 ng) was oxidized by incubating with 16 micrograms of TET2 (EM-seq TET2 Reaction Buffer (reconstituted), E7128A, E7131A diluted and E7130A) for 30 min at 37°C followed 30 min incubation with BGT (EM-seq component E7129A, NEB, Ipswich, MA, USA) in the same buffer at 37°C. Oxidized brain genomic DNA was incubated additional 30 min with 0.8U of Proteinase K at 37°C, and subsequently purified with Genomic DNA Clean Concentrator kit (Zymo Research, Irvine, CA, USA Research). Purified DNA was denaturated at 80°C in presence of 66% of formamide, and deaminated with 0.3 micrograms of APOBEC3A in 100 ml reaction volume (EM-seq components E7133AA and E7134AA) for 16 hours for 5-hmC detection and 3 hours for 5-mC detection. We then purified DNA with Genomic DNA Clean Concentrator kit (Zymo Research, Irvine, CA, USA Research). Targeted DMRs were amplified from each of the purified deaminated DNA using custom designed primers (Supplemental Table 6).

#### Single Molecule Real Time (SMRT) sequencing

The purified long amplicons were prepared for PacBio SMRT sequencing (Pacific Biosciences, Menlo Park, CA, USA) following the “Amplicon template preparation and sequencing” protocol. One library was prepared for each region and for each modification type and was loaded onto SMRT cell using the MagBead method. The LR-EM-seq libraries were sequenced on a PacBio RSII machine with 5.5-hour movie. Consensus sequences of individual sequenced molecules (Read of Insert) were generated by the “RS ReadsOfInsert” protocol using the SMRT portal (Version 2.3.0.140893). Reads that were shorter than the expected amplicon size were removed from downstream analysis. We then corrected the quality scores of the SMRT consensus sequences using the BBmap tools (Bushnell B. - sourceforge.net/projects/bbmap/) and then conducted phasing analysis (see below “5-mC and 5-hmC phasing analysis”).

### Bisulfite conversion for the damage assay

200 ng of mouse E14 genomic DNA sheared to 15 kb fragments were treated with sodium bisulfite using an EZ DNA Methylation-Gold Kit (Zymo Research, Irvine, CA, USA Research), or an EpiTect Bisulfite Kit (QIAGEN, Hilden, Germany), according to the manufacturers’ recommendations. Primers used to amplify bisulfite converted DNA are listed in Supplementary Table 7.

### Whole genome library preparation and sequencing

#### 5-mC

50 ng mouse E14 genomic DNA, spiked with 0.5% unmethylated Lambda DNA (Promega, Madison, WI, USA), was sheared to 250 bp fragments with a Covaris S2 sonicator (Covaris). Fragmented DNA was used for library preparation using a NEBNext Ultra II Kit (NEB, Ipswich, MA USA) according to the manufacturer’s instructions for DNA end repair, methylated adapter ligation, and size selection. The adapter ligated DNA fragments were deaminated by the enzymatic deamination method using Enzymatic Methyl-seq Conversion Module (NEB, E7125) or by sodium bisulfite using two commercially available kits: BS (1)EZ DNA Methylation-Gold Kit (Zymo Research, Irvine, CA, USA Research) and BS (2) EpiTect Bisulfite Kit (QIAGEN, Hilden, Germany) respectively according to the manufacturers’ recommendations. The deaminated DNA were amplified with Q5 dU Bypass DNA polymerase Master Mix (NEB, Ipswich MA, USA) and the resulting libraries were analyzed and quantified with an Agilent Bioanalyzer 2100 DNA High-sensitivity chip. All the whole-genome libraries were sequenced using the Illumina NextSeq platform with 25% phiX spike-in. Pair-end sequencing of 150 cycles (2 × 150 bp) was performed for all the sequencing runs. Base calling and demultiplexing were carried out with the standard Illumina pipeline.

#### 5-hmC

50 ng of mouse E14 DNA, spiked with 0.5% of methylation free lambda (Promega, Madison, WI, USA), T4gt and Xp12 phages DNA, was sheared with a Covaris S2 instrument using the recommended settings for 250 bp fragments. Sonicated DNA was then incubated with 20U of BGT (NEB) in 57 ml using end repair buffer from NEBNext Ultra II DNA library preparation kit for Illumina (NEB). Glucosylated DNA was then end repaired without purification followed by ligation to Pyrrolo-dC adaptors as indicated in NEBNext Ultra II DNA library preparation protocol (NEB, Ipswich, MA, USA) with the following modification to the protocol: The Antarctic USER (M0507, NEB, Ipswich, MA, USA) was used instead of USER to open up Pyrrolo-dC loop adaptor. DNA was SPRI bead purified, denaturated at 80°C in presence of 66% of formamide, and deaminated with 0.3 µg of APOBEC3A (EM-seq component E7133A) in 100 µl reaction volume (1x reaction buffer EM-seq component E7134A) for 16 hours. After SPRI bead purification, the library was amplified with Q5 Bypass U DNA polymerase (M0598, NEB, Ipswich MA, USA). The amplified libraries were purified and sequenced using Illumina NextSeq 500 platform with paired-end sequencing (2 × 150bp).

#### Global nucleoside analysis

Genomic control DNAs were digested to nucleosides by treatment with the Nucleoside Digestion Mix (NEB, M0649S) for 1 h at 37 C. LC-MS analysis was performed on an Agilent LC/MS System 1200 Series equipped with a G1315D diode array detector and a 6120 Single Quadrupole Mass Detector operating in positive (+ESI) and negative (-ESI) electrospray ionization modes. LC was carried out on a Waters Atlantis T3 column (4.6 150 mm, 3 µm) with a gradient mobile phase consisting of 10 mM aqueous ammonium acetate (pH 4.5) and methanol. The relative abundance of each nucleoside was determined by UV absorbance. LC-MS/MS analysis was performed in duplicate by injecting digested DNA on an Agilent 1290 UHPLC equipped with a G4212A diode array detector and a 6490A Triple Quadrupole Mass Detector operating in the positive electrospray ionization mode (+ESI). UHPLC was carried out on a Waters XSelect HSS T3 XP column (2.1 100 mm, 2.5 µm) with the gradient mobile phase consisting of methanol and 10 mM aqueous ammonium formate (pH 4.4). MS data acquisition was performed in the dynamic multiple reaction monitoring (DMRM) mode. Each nucleoside was identified in the extracted chromatogram associated with its specific MS/MS transition: dC at m/z 228→112, 5-mC at m/z 242→126, and 5-hmC at m/z 258→142.. External calibration curves with known amounts of the nucleosides were used to calculate their ratios within the samples analyzed

### Bioinformatics analysis

#### data processing and 5-mC, 5-hmC calling of whole genome sequencing libraries

Raw reads were first trimmed by the Trim Galore software (https://github.com/FelixKrueger/TrimGalore) to remove adapter sequences and low-quality bases from the 3’ end. Unpaired reads due to adapter/quality trimming were also removed during this process. The trimmed read sequences were C to T converted and were then mapped to a composite reference sequence including the mouse genome (mm9) and the complete sequences of lambda, pUC19, phage XP12 and T4 controls using the Bismark program [23] with default Bowtie2 setting. The aligned reads were then subjected to two post-processing QC steps: 1, alignment pairs that shared the same alignment start positions (5’ ends) were regarded as PCR duplicates and were discarded; 2, reads that aligned to the mouse genome and contained excessive cytosines in non-CpG context (e.g., more than 5 in 150bp) were removed because they likely resulted from conversion errors. The de-duplication step was skipped for loci-specific amplicon libraries. The remaining good quality alignments were then used for cytosine methylation and hydroxymethylation calling by Bismark methylation extractor.

#### Statistical inference of high-confident methylated and hydroxymethylated cytosines

We applied a binomial distribution B (n,p) [24] to identify 5-mC and 5-hmC sites with < 1% false discovery rate. In this binomial distribution, the number of trials (n) is the sequencing depth at each cytosine position. The error rate corresponds to the probability p in the binomial distribution and was estimated from the unmethylated Lambda genome. The error rates were calculated from unmethylated lambda genomic DNA (EM-seq (5-mC) error rate: 0.001 for both CpG and CpH; Bisulfite method (1) error rate: 0.016 for CpG and 0.015 for CpH; Bisulfite method (2) error rate: 0.004 for both CpG and CpH). For 5-hmC detection libraries made using APOBEC(5hmC) method, the non-conversion error rate of unmodified cytosines is estimated from unmethylated lambda DNA and was 0.001 for both CpG and CpH. The non-conversion error rate of methylated cytosines was estimated from fully methylated XP12 genomic DNA and was 0.01 for all both CpG and CpH.

#### Genotype correction for methylated cytosines

Because the mouse embryonic stem cells used in this study have different genotypes from the mouse reference sequence (mm9), we examined the sequencing data of the +1 position of every identified methylated cytosines in non-CpG context (w.r.t the reference sequence) to determine whether the base is a true H or a SNP of G in our cell line. If the sequenced bases of the +1 position from both strands consistently indicate guanine (i.e., the corresponding bases on the opposite strand were either cytosine or thymine – converted from unmethylated cytosine), we then correct the methylation context to mCpG. If the +1 bases were heterozygous for a SNP, we then discarded them from the analysis.

#### Correlation analysis of read coverage and cytosine content

We first removed all the PCR duplicates and used Read 1 of aligned read pairs with high mapping quality (MAPQ>20) for this analysis. We calculated the number of the original cytosines and the number of converted cytosines for each aligned read. The distribution of reads with respect to bins of different cytosine content (cytosine content is calculated as number of Cs/alignment length and is divided into 20 bins at 5% intervals) or bins of different density of converted cytosines (density of converted cytosines is calculated as number of converted Cs/ alignment length) was calculated for each library. And the distribution was further normalized to the distribution of 100bp windows across the entire reference genome (mm9) with respect to background cytosine content.

#### Correlating 5-mC and 5-hmC to ChIP-seq Data Sets

All the external ChIP-seq data sets were downloaded from the NCBI GEO database (TET1: GSE24843 [25]; RNA polymerase II: GSE12241 [26],Transcription factors: GSE11431 [27]. Histone modification marks H3K4me3: GSE12241 [26]and H3K4me1: GSE24165 [28]. For TET1, RNA polymerase II and histone marks, we downloaded the mapped reads of the ChIP-seq experiments and used the MACS2 program [29]to identify peaks of binding sites of each data set (Tet1: peak p value < 10^−8^, fold enrichment over IgG > 10; RNA polymerase II: peak p value < 10^−5^, fold enrichment over control > 10; histone marks: peak p value < 10^−5^, fold enrichment over control H3 > 10). For datasets that were originally mapped to the mm8 reference genome (RNA polymerase II and histone modification marks) we remapped them the mm9 reference using the LiftOver tool [30] prior to the MACS2 peak calling analysis. For the 13 transcription factors we used the predicted binding sites directly and remapped the genomic coordinates to the mm9 reference using the LiftOver tool [30]. 5-mC and 5-hmC sites were mapped to individual regions using BEDTOOL [31]. Site density was calculated as number of sites divided by region length. 5-mC and 5-hmC densities was also normalized by background cytosine and 5-mC densities respectively. Average densities were then computed for each bin position and were used for meta-plots or global trend plots.

#### 5-mC and 5-hmC phasing analysis

We used Bismark [23] to map full-length reads from Pacbio SMRT sequencing and Oxford Nanopore sequencing to the mouse reference genome (mm9) with the following parameters: –bowtie2 -N1 -L15 –score min L,0,-0.6. The modification states of individual CpG sites were called by the bismark methylation extrator program. We then extracted the context specific methylation information of individual molecules and plotted in R. SNPs were called using the SAMtools [32]. We used the SNPs that were also reported previously as heterozygous SNPs [33] to distinguish paternal and maternal copies in the F1 sample for phasing analysis. The conversion rates were calculated using all the cytosines in CpC and CpT context by formula: converted C(C/T)/ total C(C/T). We exclude CpA from the calculation of non-conversion error rate because it was previously reported that brain has high level of CpA modification [34] [33] [35].

## Acknowledgments

We thank Yi Zhang and Hao Wu for DNMT TKO J1 ES cell genomic DNA, Peter Weigele for providing XP12 phage DNA, Laurie Mazzola, Joanna Bybee and Danielle Rivizzigno for sequencing.

