## Supplementary Text and Figures for "Non-destructive enzymatic deamination enables single molecule long read sequencing for the determination of 5-methylcytosine and 5-hydroxymethylcytosine at single base resolution"

**Supplementary text 1: Validation of the EM-seq technology**

To validate EM-seq in the identification of 5-mC, we used control samples consisting of 100 ng mouse embryonic stem cell (E14) genomic DNA spiked with 0.5% methylation free lambda DNA. We performed both enzymatic deamination and bisulfite conversion using two widely used bisulfite kit using the same starting material (**Material and Methods**). Technical replicates were also performed to assess reproducibility of the methods.

The resulting libraries were sequenced using Illumina platform and down-sampled to ~118 Million reads per library. The resulting paired-end reads were mapped to the mouse genome (mm9) and the phage lambda genome (**Material and Methods**). From the unmethylated phage lambda, enzymatic deamination experiments show a non-conversion error rates of 0.1% (**Supplementary table S1**) with no apparent sequence preference (**Supplementary Fig. 1A**). In comparison, the non-conversion error rates of bisulfite treated lambda DNA are ranging from 0.4 % to 1.6 % depending on the BS kit used, which is in line with the usual non-conversion rates reported in the literature for whole genome bisulfite sequencing (WGBS) (**Supplementary table S1**). Importantly, for the bisulfite treated DNA, we observed a higher level of residual unconverted cytosines in CpA context. This CpA bias is also observed with the published WGBS dataset (GSM432685)^1^ (**Supplementary Fig. 1B**). This bias leads to 10 to 40-fold more false positive high-confident methylated CpA in WGBS compared to enzymatic conversion (**Supplementary Fig. 1C**). This overestimation of the amount of methylated CpA in bisulfite treatment is anticipated to directly confound the identification of methylation in non-CpG context. This may alter the interpretation of epigenetic marks in notably CpA context, a predominant non-CpG methylation recently reported in embryonic stem cells ^1^ ^2^ ^3^ ^4^, brain ^5^ ^6^ ^7^ and germ cells ^8^.

Identification of cytosine methylation in the mouse DNA reveals consistent results between enzymatic deamination and WGBS with a characteristic bimodal distribution of genome-wide methylation abundance in CpG context (**Supplementary Fig. 1D**). CpG methylation calls between enzymatic conversion and bisulfite sequencing are ~96% agreement (**Supplementary Fig. 1E**). Methylation calls from enzymatically treated DNA were highly reproducible between technical replicates (**Supplementary Fig 1F**) indicating robustness and reproducibility of the methodology. Furthermore, CpG methylation profiles in mouse stem cells revealed using enzymatic conversion are correlated with markers of repressive and active chromatin with 5-mC depletion in active transcription regions and promoters (**Supplementary Fig. 1G & 1J**).

Genome-wide investigation of the read distribution in the mouse genome reveals that the enzymatic deamination method produces more even sequencing coverage than the bisulfite conversion based sequencing methods (**Supplemental Fig. 2A**) and consequently yields more useful data. For example, at equal amount of sequencing reads (118M) and considering only sites with at least 5X coverage (minimal coverage for methylation calling), enzymatic conversion covers 79 % of all CpG pairs compared to 73 % of CpG pairs in WGBS. This difference corresponds to 1.2 million additional CpG pairs in the enzymatic conversion method as a result of a more uniform genome-wide sequencing coverage compared to both WGBS. More uniform coverage from enzymatic deamination results in a notable increase in coverage notably in CpG rich regions of the genome such as promoters and CpG islands (**Supplementary Fig. 2B**). We reasoned that bisulfite-induced damage at cytosines can result in a bias in coverage for WGBS and investigated the relationship between read coverage and cytosine density. Supporting large scale damage at cytosine previously reported ^9^, coverage of WGBS is inversely correlated with cytosine and deaminated cytosines densities. Consequently, read mapping to regions containing fewer cytosines/converted cytosines are over-represented (**Supplementary Fig. 2C and supplemental Fig. 2D)**.  On the other hand, enzymatic conversion shows more uniform coverage across wide ranges of cytosine contents and was consequently able to cover a higher fraction of CpG pairs, notably in cytosine rich regions. Consistent with the non-destructive nature of enzymatic reaction, these results indicate that enzymatic conversion preserves the integrity of the DNA.

**Supplementary text 2: Simultaneous analysis of 5-mC and 5-hmC in mouse embryonic stem cell**

We Conducted both enzymatic 5-mC and 5-hmC sequencing in parallel to enabled for the first time to simultaneously investigate 5-mC and 5-hmC using the same baseline enzymatic reaction. We first explored the relative levels of 5-hmC and 5-mC in TET1 binding sites previously defined in mouse E14 cell line using ChIP-seq ^10^. Because TET1 is a major contributor in the oxidation of 5-mC to 5-hmC ^11^, an increased level of 5-hmC is expected at TET1 binding sites. As expected, we observed a decrease of 5-mC and a local enrichment of 5-hmC at the binding site of the TET1 protein (**Supplementary** **Fig. 1J**), which is consistent with the activity of the enzyme. We also confirmed previous observation ^12^ that RNA polymerase II (Pol II) binding regions have low levels of both 5-mC and 5-hmC (**Supplementary** **Fig. 1J**) in agreement with a common understanding that core promoters of actively transcribed regions are in general depleted of cytosine modifications.

Next, we analyzed the cytosine modifications around the binding site of key transcription factors. We found that, while 5-mC are globally depleted in these regions, the profile of hydroxymethylation varies largely according to the type of transcription factor. Most of the transcription factors (TFs) showed increased enrichment in hydroxymethylation. Such an enrichment can be directly at the binding site (Tcfcp2l1, Stat3 and Esrrb) or is bimodal around the binding site (Nanog, Oct4, Sox2 and Klf4), suggesting that the deposition of 5-hmC is not only related to the binding of transcription factors, but also affected by other factors. Strengthening this observation, we also observed that distal enhancers marked by H3K4me1 and the absence of H3K4me3 displayed similar enhanced 5-hmC levels. However, the extend of 5-mC depletion in the enhancer regions is less pronounced than that in the TF binding *loci* implying that 5-hmC is not simply an intermediate of DNA demethylation pathway.

Some transcription factors showed no significant changes or slight depletion of 5-hmC compared with the corresponding flanking regions (E2f1, Zfx, c-Myc and n-Myc) in agreement with their tendency to predominantly bind to core-promoter regions. Previous studies have reported depleted DNA methylation at DNA-protein interaction sites ^1^ ^13^ and enrichment of hydroxymethylation near key transcription factor binding sites in mouse embryonic stem cells ^14^. In agreement with these studies, we detected local depletion of DNA methylation at the binding sites of all the important transcription factors. Moreover, we observed lower than genome average 5-mC abundance in the adjacent regions beyond the transcription factor footprints. This result implies that the depletion of DNA methylation may be primed upon other activities that affect the DNA more extensively, such as changes of chromatin structure.

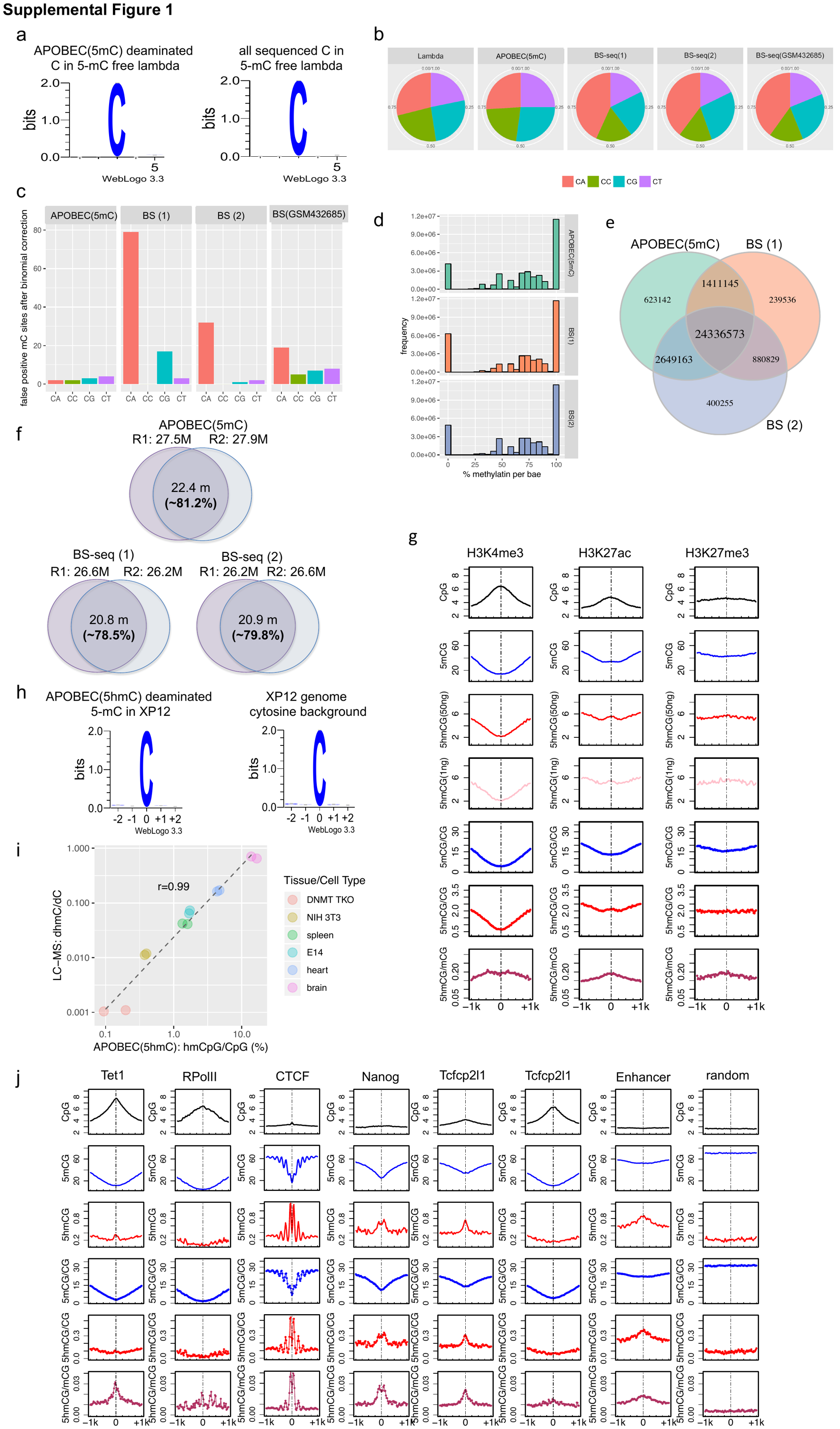

**Supplementary Figure 1**: **a.** Deaminated cytosines in the unmethylated lambda genome showed no systematic sequence preference by APOBEC(5mC)-based deamination method. **b.** Pie charts show the proportions of the four cytosine dinucleotides sequence context (CpA, CpC, CpG and CpT) of the erroneous methylation calls in the unmethylated lambda genome from four whole-genome sequencing experiments (APOBEC(5mC), two whole-genome BS-seq i.e., BS-seq (1) and BS-seq (2),  and one published whole-genome BS-seq experiment (GSM432685)) in comparison to the lambda genomic composition (left). **c.** Numbers of false positive 5-mC sites in the unmethylated Lambda control in each dinucleotide sequence context after binomial correction using library specific average non-conversion error rate. All the BS-seq libraries including a public dataset (GSM432685) had significantly more false positive 5-mC sites in the CpA context than other contexts. **d.** Histograms represent genome-wide CpG methylation levels from 0% to 100% distributed across 20 bins of 5% intervals for EM-seq and 2 WGBS libraries.  **e.** Comparison of CpG sites between enzymatic deamination method (APOBEC(5mC)) and WGBS. 95.8% and 95.5% of the total identified 5-mCpG sites by the two BS-seq libraries respectively are also identified by enzymatic deamination method. Three font sizes are used as indicators for the relative magnitude of the numbers. The large font size is used for numbers greater than 10 million, the middle front size is used for numbers whole values are between 1 million and 10 million; and the small font size is used for numbers smaller than 1 million. **f.** Overlap of called 5-mCpG sites between the two technical replicates of the 3 protocols: enzymatic deamination method (APOBEC(5mC)), BS-seq (1) and BS-seq (2). **g.** Distribution patterns of 5-mCpG (blue), 5-hmCpG (red: 50 ng library; pink: 1 ng library) and normalized levels to background CpG density at active transcription chromatin mark (H3K4me3), active enhancer mark (H3K27ac) and repressive chromatin mark (H3K27me3). **h.** Deaminated 5-mC in the fully methylated XP12 genome showed no systematic sequence preference by APOBEC(5hmC) enzymatic deamination method. **i.** Pearson correlation between 5-hmC measured using sequencing of enzymatic deaminated DNA (x-axis) versus LC-MS (y-axis) for various genomic DNA. There are two technical replicates of each sample. **j.** Distribution patterns of 5-hmC (red) and 5-mC (blue) at protein-DNA pinding sites. The absolute (smooth lines) and normalized (dotted lines) 5-hmC (50 ng library) and 5-mC levels in the CpG context are depicted around Tet1 and RNA polymerase II binding sites as well as ESC-related transcription factor binding sites and enhancer regions. Unbound sites which are random sampled from the reference genome served as control.

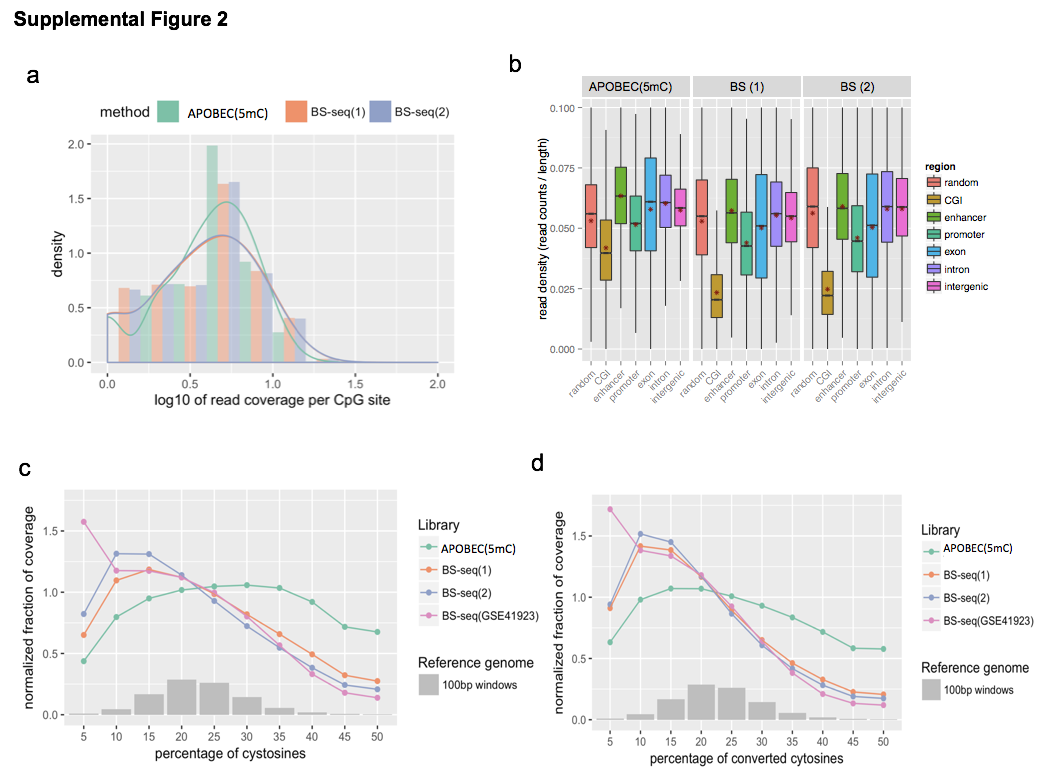

**Supplementary Figure 2: a.** Distribution of CpG sequencing copies of the APOBEC enzymatic deamination library vs. the two whole-genome BS-seq libraries, i.e., BS-seq (1) and BS-seq (2). **b.** Read coverage at different genomic regions for the enzymatic deamination method (APOBEC(5mC), left panel) and two bisulfite conversion methods (BS(1) and BS(2), right panels). At equal number of reads, the enzymatic deamination method yields notably higher coverage in cytosine-rich regions such as promoter and CpG islands leading to a more even coverage across features compared to both bisulfite conversion methods. **c.** Relationship between read coverage and cytosine density: colored lines represent the normalized proportions of read coverage at various cytosine content bins (normalized coverage is calculated as proportion of reads at a specific C% bin divided by fraction of the 100-bp windows across the reference genome, which is illustrated by the histogram in grey). The enzymatic deamination method APOBEC(5mC) (green) shows the least coverage bias relative to cytosine content and covers notably more cytosine rich regions compared to three bisulfite sequencing libraries including a published bisulfite dataset (purple). **d.** Relationship between read coverage and density of converted cytosines: colored lines represent the normalized proportions of read coverage at various densities of converted cytosines (normalized coverage is calculated as proportion of reads at a specific bin of converted cytosine density divided by fraction of the 100-bp windows across the reference genome, which is illustrated by the histogram in grey). The library by enzymatic deamination method (green) has the least biased sequencing coverage in terms of converted cytosine content.

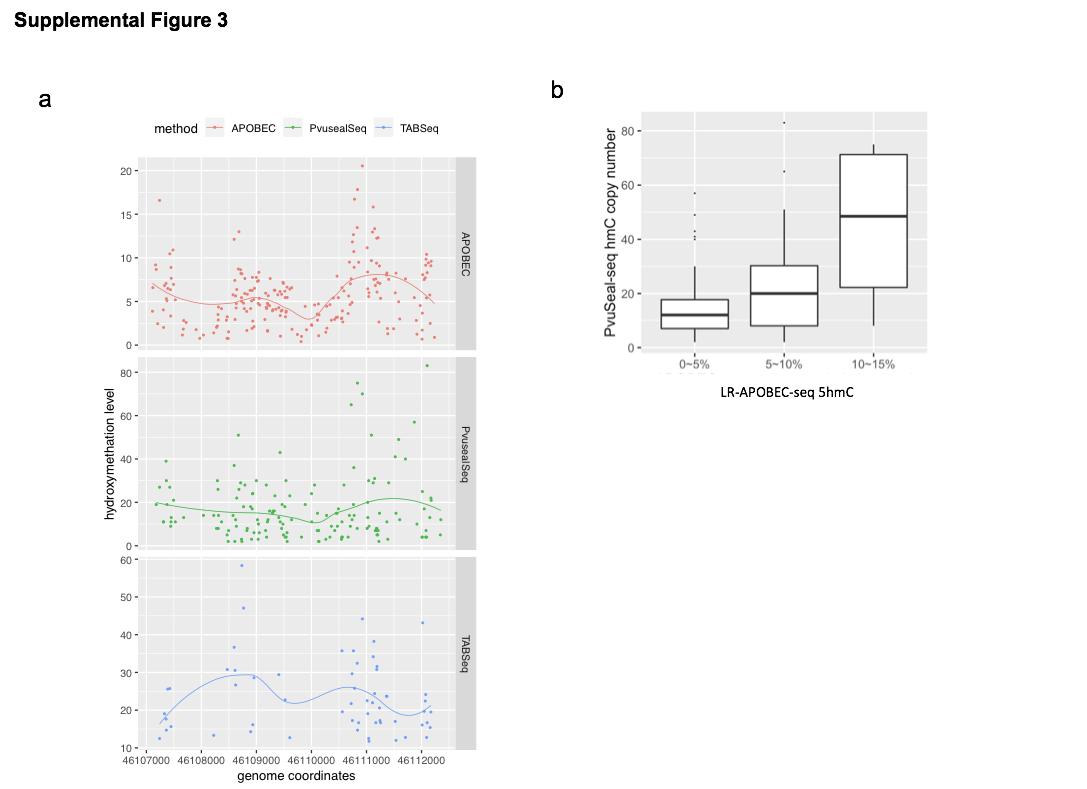

**Supplemental Figure 3: a.** Single-base hydroxymethylation level of CpG sites within the 5078 bp region from the mouse E14 cell calculated from 3 methods: LR-EM-seq (top), PvuSeal-seq (middle) and TAB-seq (bottom). The fitted lines are drawn using the LOESS method. In the cases of LR-EM-seq and TAB-seq, the y-axis shows the percentage of hydroxy-methylated copies among all the sequenced copies (modified + unmodified). In the case of Pvu-Seal-seq, the y-axis shows the number of captured modified copies, which is a relative measurement of hydroxymethylation level. **b.** LR-EM-seq hydroxymethylation measurements are in good agreement with PvuSeal-seq results. CpG sites were divided into 3 subgroups based on LR-EM-seq measured hydroxymethylation level. For each subgroup, hydroxymethylation level reported by Pvu-Seal-seq method (represented by sequencing copy number as a relative quantification) was depicted by boxplot.

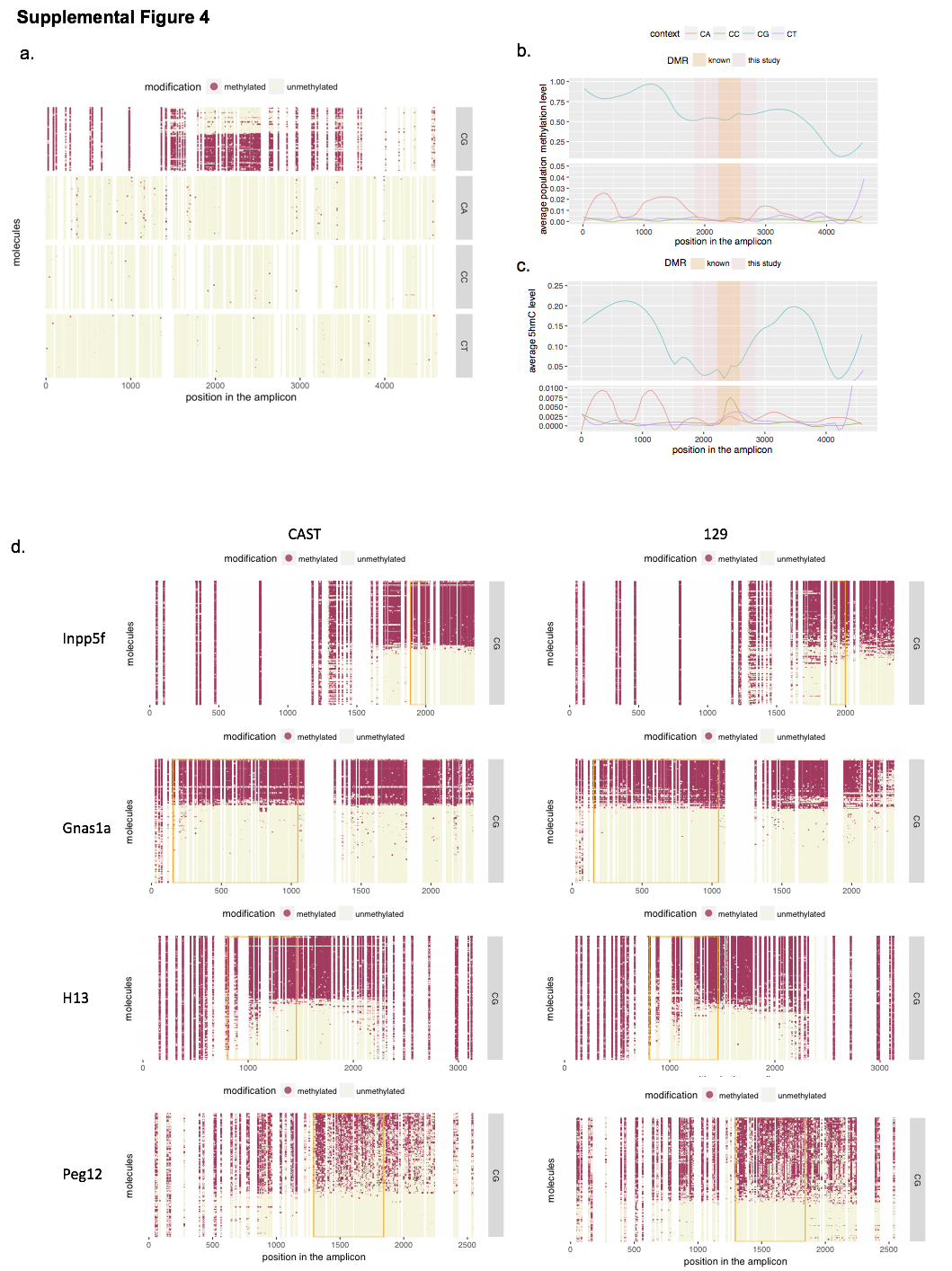

**Supplemental Figure 4: a.** Methylation states (beige: unmethylated; marron: methylated) of individual cytosines at each dinucleotide context (CpG, CpA, CpC, CpT) at single molecule level of the 4.6 kb region of the imprinted *Inpp5f_v2* gene. CpA methylation (2nd^d^ panel) is completely missing from the imprinted DMR (top panel). **b.** Trend plot shows the population level methylation abundance of each cytosine dinucleotide context in the amplified region of *Inpp5f_v2* gene. CpA methylation (red) appears to be depleted in the DMR. **c.**  Trend plot shows the population level hydroxy-methylation abundance of each cytosine dinucleotide context in the amplified region of *Inpp5f_v2* gene. **d.** Methylation states (beige: unmethylated; marron: methylated) of individual CpG sites at single molecule level of 4 previously reported DMR regions from two mouse inbred strains: Cast/EiJ (left) and 129X1/SvJ (right).

**Supplementary Table 1. Conversion rates in LR-APOBEC-seq, bisulfite sequencing and published bisulfite sequencing studies.**

| **ID** | **Conversion Rate%** | **Method** | **Conversion Rate Estimation Method** | **Study Organism** | **Reference** |
| --- | --- | --- | --- | --- | --- |
| APOBEC | 99.9 | APOBEC (5-mC) | unmethylated lambda DNA | mouse | Present study |
| BS1 | 98.5 | BS1 | unmethylated lambda DNA | mouse | Present study |
| BS2 | 99.6 | BS2 | unmethylated lambda DNA | mouse | Present study |
| 1 | 99.3~99.8 | WGBS | unmethylated lambda DNA | human | Lister et al., 2009 ^15^ |
| 2 | 99.5 | WGBS | unmethylated lambda DNA | mouse | Xie et al., 2012 ^6^ |
| 3 | 98.5~99.6 | WGBS* | unmethylated lambda DNA | human | Consortium et al., 2015 ^16^ |
| 4 | 97.2~99.8 | WGBS | unmethylated lambda DNA | Zebrafish, Xenopus tropicalis, mouse | Bogdanovic et al., 2016 ^17^ |
| 5 | 97.7 | single cell WGBS | non-CpG sites | mouse | Smallwood et al., 2014 ^18^ |
| 6 | >99 | single cell WGBS | custom-designed unmethylated oligonucleotides | human and mouse | Farlik et al., 2015 ^19^ |
| 7 | 97.6~ 99.1 | single cell WGBS | non-CpG sites in the genome | mouse | Gravina, Dong, Yu, & Vijg, 2016 ^20^ |
| 8 | >97 | RRBS |  | mammals | Smith, Gu, Bock, Gnirke, & Meissner, 2009 ^21^ |
| 9 | >=99 | ERRBS  (enhanced RRBS) | Unmethylated C | human and other animals | Garrett-Bakelman et al., 2015 ^22^ |
| 10 | 97 | SMRT -BS | non-CpG Cytosines | human | Yang et al., 2015 ^23^ |

* This study also conducted RRBS and MeDIP-seq, but only WGBS conversion rates were reported here.

**Supplementary Table 2. Conversion rates of Enzymatic deamination, TAB-seq and ACE-seq**

| **Method** | **C**  **conversion rate** | **5-mC**  **conversion rate** | **5-hmC protection** | **Reference** |
| --- | --- | --- | --- | --- |
| **Enzymatic deamination** | 99.9 | 98.4 | 98.4 | current study |
| **TAB-seq** | 99.62 | 97.79 | 84.4~92 | Yu et al., 2012 ^24^ |
| **TAB-seq** | 99.57 | 98.47 | 57 | Bogdanovic et al., 2016 ^17^ |
| **TAB-seq** | 99.99 | 99.98 | 75 | Bogdanovic et al., 2016 ^17^ |
| **TAB-seq** | 98.3 | 99.61 | 55.91 | Bogdanovic et al., 2016 ^17^ |
| **ACE-seq** | 99.9% CHG  99.9% CHH  99.5% CpG | 97% | CHG 98.6%  CHH 98.5%  CpG 98.5% | Schutsky et al., 2018 ^25^ |

**Supplementary Table S3.**  **Hydroxymethylation quantification results for six mouse tissue/cell types by the enzymatic deamination method**

| **Cell/Tissue** | **5hmCpG%**  **(R1, R2)** | **5hmCpH%**  **(R1, R2)** |
| --- | --- | --- |
| **DNMT TKO** | 0.1, 0.2 | 0.1, 0.1 |
| **NIH/3T3** | 0.4, 0.4 | 0.1, 0.1 |
| **Spleen** | 1.5, 1.5 | 0.1, 0.1 |
| **E14** | 1.6, 1.6 | 0.1, 0.1 |
| **Heart** | 4.6, 4.6 | 0.1, 0.1 |
| **Brain** | 14.7, 14.8 | 0.2, 0.2 |

**Supplementary Table 4. Enzymatic 5-hmC library sequencing metrics**

| Starting Material | Replicate | Reads (million) | mapping efficiency | duplication level | CpG hmC% | CH hmC% | conversion rate C | conversion rate ^5m^C | BGT protection rate |
| --- | --- | --- | --- | --- | --- | --- | --- | --- | --- |
| 50ng mES gDNA | 1 | 127 | 67.5% | 4.2% | 4.6 | 0.1 | 99.9% | 98.4% | 98.4% |
| 50ng mES gDNA | 2 | 124 | 68.6% | 4.3% | 4.6 | 0.1 | 99.9% | 98.4% | 98.4% |
| 1ng mES gDNA | 1 | 165 | 69.9% | 58.9% | 4.6 | 0.2 | 99.8% | 98.5% | 96.7% |
| 1ng mES gDNA | 2 | 145 | 70.1% | 57.0% | 4.7 | 0.2 | 99.8% | 98.6% | 97.1% |

**Supplementary Table 5. Percentage of converted cytosines of various sequence contexts calculated from the SMRT sequencing reads of the 4616 bp Inpp5f DMR amplicon obtained using LR-APOBEC-seq.**

| **5-mC context** | **number of unconverted Cs** | **number of converted Cs** | **Unconverted C%** |
| --- | --- | --- | --- |
| **CpA** | 810 | 104053 | 0.008 |
| **CpC** | 106 | 81165 | 0.001 |
| **CpG** | 25653 | 20166 | 0.560 |
| **CpT** | 315 | 122910 | 0.003 |
| **CpC, CpT** | 421 | 204075 | 0.002 |
| **CpH (A,C,T)** | 1231 | 308128 | 0.004 |
| **5-hmC context** | **number of Ts** | **number of Cs** | **T/(C+T)** |
| **CpA** | 178 | 63849 | 0.003 |
| **CpC** | 56 | 49772 | 0.001 |
| **CpG** | 2122 | 25928 | 0.076 |
| **CpT** | 73 | 75623 | 0.001 |

**Supplemental Table 6: Primers used in the LR-APOBEC-seq experiments**

| Amplicon coordinates | Associated gene | Forward Primer | Reverse Primer | Used for |
| --- | --- | --- | --- | --- |
| chr7:135829750-135832151 | *Inpp5f* (imprinted) | GGAAGAAAGTGAGTTGATATTTTAGAGTTAG | CACTAACACTTTAACCATAAATCCTACAA | 129X1/SvJ and Cast/EiJ mouse brain |
| chr7:135829750-135832875 | *Inpp5f* (imprinted) | GGAAGAAAGTGAGTTGATATTTTAGAGTTAG | CATAAACTCATTCTACCAACCAATATC | F1 (129X1/SvJ male and Cast/EiJ female) brain |
| chr7:135829567-135834182 | *Inpp5f* (imprinted) | TGATATTTAGGGTTGTTAAGGTTTGA | CAACAACACTTCCACATCTCTATAC | mouse brain |
| chr2:174150371-174153639 | *Gnas1a* (imprinted) | TATAGTAATGTTGTTGTTTTTGAAAGTAGATG | ACTATTACCCTTAAATACTCCTTAAATTAACTCTT | 129X1/SvJ and Cast/EiJ mouse brain |
| chr2:174152457-174154782 | *Gnas1a* (imprinted) | AGGAAGGTGGTTTAAAATTTTTGAT | AACCCAAAAACACTCTAAAACAAC | 129X1/SvJ and Cast/EiJ mouse brain |
| chr7:69607178-69610032 | *Peg12* (imprinted) | AGGGTTAGAAGGAGGAAATTAATTT | CTAATATAATTATCCAAATTCCCATTCAA | 129X1/SvJ and Cast/EiJ mouse brain |
| chr2:152511205-152514436 | *H13* (imprinted) | AATTAAGAGGTTTTGGTTTGTTGGT | ACACCAAATCCTTAATATCAAACTCATA | 129X1/SvJ and Cast/EiJ mouse brain |
| chr8:46107176-46112553 | *FAT1* | AATGAAGGAAATGAATTTGGTAGAG | TCCCAAATACATAAATCCACACTTA | mouse embryonic stem cell |
| T4 phage  5357 bp | N/A | GCATAGTTCATTGAAGAGGATTTAATAG | CAACAAAACGATAAAAAGTTGTTTAC | T4 phage  Control for 5-hmC protection |
| Lambda phage  3233 bp | N/A | TAGTGAGGTAGATTTTTAGTTAGGAATTATTG | TACATATTCTACAACATATCAAACATCTTCAT | Lambda phage  Control for C deamination rates |
| CpG methylated pUC19  1774 bp | N/A | GGAAGTATAAAGTGTAAAGTTTGGGGTG | ATTTTCCAATAATAAACACTTTTAAAATTC | CpG methylated pUC19  Control for 5-mC protection |

**Supplemental Table 7: Primers for DNA damage assay**

| Primers for Integrity assessment (Fig. 2e) | | |
| --- | --- | --- |
| Amplicon size (bp) | Forward primer | Reverse primer |
| 388 | TAGGATAAAAATATAAATGTATTGTGGGATGAGG | AAAACATATAACCCCCTCCACTAATAC |
| 731 | AGATATATTGGAGAAGTTTTGGATGATTTGG | AAAACATATAACCCCCTCCACTAATAC |
| 1456 | TAAGATTAAGGTAGGTTGGATTTGG | TCATTACTCCCTCTCCAAAAATTAC |
| 2018 | AAGATTTAAGGGAAGGTTGAATAGG | ACCTACAAAACCTTACAAACATAAC |
| 3325 | TGGAGTTTGTTGGGGGGTTTGTTGTTTAAG | TCTAACCCTCACCACCTTCCTAATACCCAA |
| 4229 | TGGTAAAGGTTAAGAAGGGAAGATTGTGGA | AACCCTACTTCCCCCTAACAAATTTTCAAC |
| Primers for qPCR assay (Fig. 2c) | | |
| 336 | AGATATATTGGAGAAGTTTTGGATGATTTGG | TCCTTCCTACCCCTAACCTCTACTA |
| 456 | TTTATAGGTTGTTTGTTGTTGGAATTTTTTTTG | TCCTTCCTACCCCTAACCTCTACTA |
| 577 | AGATTAAGGTAGGTTGGATTTGGGGATTTAAG | TCCTTCCTACCCCTAACCTCTACTA |
| 809 | TTGGGTTTAGTGATTATTGGGAGGTTAG | TCCTTCCTACCCCTAACCTCTACTA |
